# Non-invasive longitudinal imaging reveals aging-associated changes in neuroimmune cells in adult zebrafish

**DOI:** 10.1101/2025.09.23.676912

**Authors:** Elizabeth M. Haynes, Corey Steinhauser, Robert Swader, George Petry, Tyler K. Ulland, Kevin W. Eliceiri

## Abstract

Zebrafish are a powerful model for imaging studies of development and neurobiology. However, most studies have focused on developing zebrafish due to technical and biological challenges of imaging adult stages. These include increased tissue opacity and illumination depth limitations, and difficulty maintaining life support and anesthesia in a 2-4cm long fish. There are currently limited tools for intravital imaging of the adult zebrafish brain. The ability to image the brain in the same individual repeatedly without physical damage would allow zebrafish to be better utilized to study aging and neurodegenerative disease. We designed and applied a 3D-printable device for non-invasive, repeatable multiphoton neural imaging of genetically non-pigmented adult zebrafish from 2 months to 19 months old. Animals successfully recovered after multi-hour imaging sessions and can be imaged repeatedly over periods of weeks to years. We show the utility of this approach through imaging the neuroimmune system, revealing that microglia in aged zebrafish have enhanced cellular dynamics. This technique could be widely used and beneficial for other cell-scale neuroimaging studies in the adult fish.

**Summary Statement:** An open-source, 3D printable device that enables non-invasive neuroimaging in adult zebrafish reveals aging-associated changes in neuroimmune cell behavior.

## Introduction

As we discover the expansive roles of microglia in the brain, there is increased interest in studying microglial biology during aging and disease progression in relevant animal models. Intravital imaging of dynamic behaviors of microglia is difficult in mammalian models and generally requires surgery or skull thinning. These procedures require a large skill and time investment and can alter the local environment of the brain(Cramer et al., 2021; Kalmbach and Waters, 2012). Small thinned skulled vertebrate teleost fish such as zebrafish and danionella are an excellent alternative model for the study of longitudinal, dynamic neuroimmune biology. In this work we describe a method to perform repeated imaging sessions over periods of weeks to months with minimal physical and behavioral stress on the animal. Several animal health concerns limit the ability to serially image individual animals, including physical trauma from mounting and unmounting, hypoxia from inadequate oxygenation, and the possibility of neurological effects from repeated sedation. To avoid some of these complications we endeavored to design an open-source 3D printable device that allows for quick and reproducible positioning of a fish in an upright position with minimal handling and use of force. Our device’s design allows the fish’s head to remain accessible to a water dipping objective in an upright orientation and keeps the gills free from any obstruction. Intubation of fish has been previously demonstrated (Barbosa et al., 2016; Castranova et al., 2022; Chow et al., 2020; Dray et al., 2015) to allow sustained oxygenation, sedation, and long-term imaging. Thus, our design is compatible with intubation as well as circulation of additional water to provide adequate water temperature control (in the range of 26-29 °C) within the chamber. Additionally, the design needed to accommodate adult fish across age ranges and length scales to facilitate longitudinal imaging in a continuously growing animal. No currently published teleost imaging device was appropriate for our specific applications (see discussion below), so this novel design contributes a valuable, easily adopted, and adaptable tool for the zebrafish research community.

There have been a handful of published approaches for imaging juvenile and adult Danio species. While zebrafish are still the most prevalent model, *Danionella* are becoming popular due to their small size, optical accessibility, and diverse behaviors including vocalization(Lam, 2022; Schulze et al., 2018; Tatarsky et al., 2022). Limited work has been done showing the ability to image other species, such as cave fish (Castranova *et al*., 2022).

Previous approaches have varied greatly in their mounting strategy, achieved depth, and goals for imaging. All approaches have used some form of intubation to provide for sedation and oxygenation. Some designs modified existing lab supplies to create a temporary mounting device (Barbosa et al., 2016, 2015; Chow et al., 2020; Dray et al., 2015). The drawback to these approaches is a lack of reproducibility both between individual experiments and across different labs, though this work demonstrated that multiphoton microscopy could be used to image brain regions of both adult zebrafish and *Danionella*.

In 2022, Castranova et al. created the first reproducible imaging chamber for studying wound healing in live adult zebrafish using an inverted microscope (Castranova *et al*., 2022). The study demonstrated that zebrafish could be imaged while under anesthesia for 20 hours while still being recoverable.

Lam et.al subsequently demonstrated a reproducible imaging chamber for *D. cerebrum* that is compatible with both upright and inverted imaging. Unlike previous examples, Lam did not rely on anesthesia to prevent movement and instead secured the fish in place by encasing it in low melting point agarose, though intubation is still used to provide oxygenation. Fish could be imaged up to 3 hours using this method. Conveniently, the smaller size of Danionella and their lack of a skull roof plate allows for high resolution neural imaging with just a standard confocal.

Development of re-usable, reproducible chambers that maintain animal viability over long period of time and repeated imaging sessions is the first step in increasing adoption of adult intravital imaging in fish species. Unfortunately, none of these strategies was highly suited to the needs of our research. Namely, we wanted a design that was non-invasive, allowed for highly reproducible upright positioning, and could maintain the health of the fish during both long-term imaging and serial imaging. We built upon these existing strategies to make an easy-to-use open-source 3D printable imaging chamber for adult zebrafish neural imaging at the cell-scale using multiphoton microscopy. Here we demonstrate its utility for non-invasive neuroimaging over hours to months.

## Results

### Non-invasive imaging of microglia distribution and dynamics in living zebrafish

To confirm the utility of our designed device (Fig. 1A, B, Sup. Fig. 2) for long-term neuroimaging, we tested the ability to image cells of the neuroimmune system in the zebrafish telencephalon and optic tectum (Fig. 1C). These areas are superficially positioned and accessible to imaging. We imaged transgenic zebrafish expressing a marker of the vasculature (*kdrl*:mCherry) along with a marker of phagocytic immune cells (*mpeg1*:eGFP) in a pigmentless Casper background (White *et al*., 2008). The inclusion of a secondary structural marker such as the vasculature is helpful to correctly identify the same regions over longitudinal imaging periods, especially since the zebrafish brain is capable of continuous neuroproliferation and growth throughout the fish’s lifespan. However, autofluorescent or second-harmonic signal in the bone or other tissues may also be suitable for re-identifying regions across time (Dray *et al*., 2015). Using two Bruker multiphoton microscopes setups, a Bruker Ultima and Ultima 2P+, we were able to image at depths of up to 500µM using a Spectra Physics Insight ultrafast laser. Laser power measured at the imaging plane at generally used settings was 10-13mW. The depth we measured from the top of the skin to the start of brain tissue varied by age and brain region, ranging from as little as 25 µM in the telencephalon of a 3-month-old fish to 225µM in the telencephalon of a 19-month old fish. We found the optic tectum tends to remain closer to the skull and thus can be easier to image in older animals. We were able to visualize both the vasculature of the telencephalon and neuroimmune cells through the intact skin and skull of adult zebrafish (Fig. 1 D, D’, Supplemental Movie 1). *Mpeg1*:eGFP positive neuroimmune cells ranged in morphology from small, rounded vessel-associated cells to larger and more ramified cells that could be close to vessels or in the parenchyma. The population of neuroimmune cells that could be imaged at laser powers generally accepted to be non-destructive tended to be superficial and just below what we presume to be the meninges, generally within the first 25-50μms of the brain tissue, but bright cells could be identified as deep as 200μm within the brain tissue. We focused on the larger, deeper, and more ramified cells as being more consistent with morphology associated with microglia. We achieved sufficient resolution to be able to identify individual cells of interest to target for imaging at greater magnification (Fig. 1E, E’, arrowheads). Cellular behavior was live imaged for 10-30 minutes for analysis (Fig. 1F, F’, Supplemental Movie 2).

**Figure 1.**
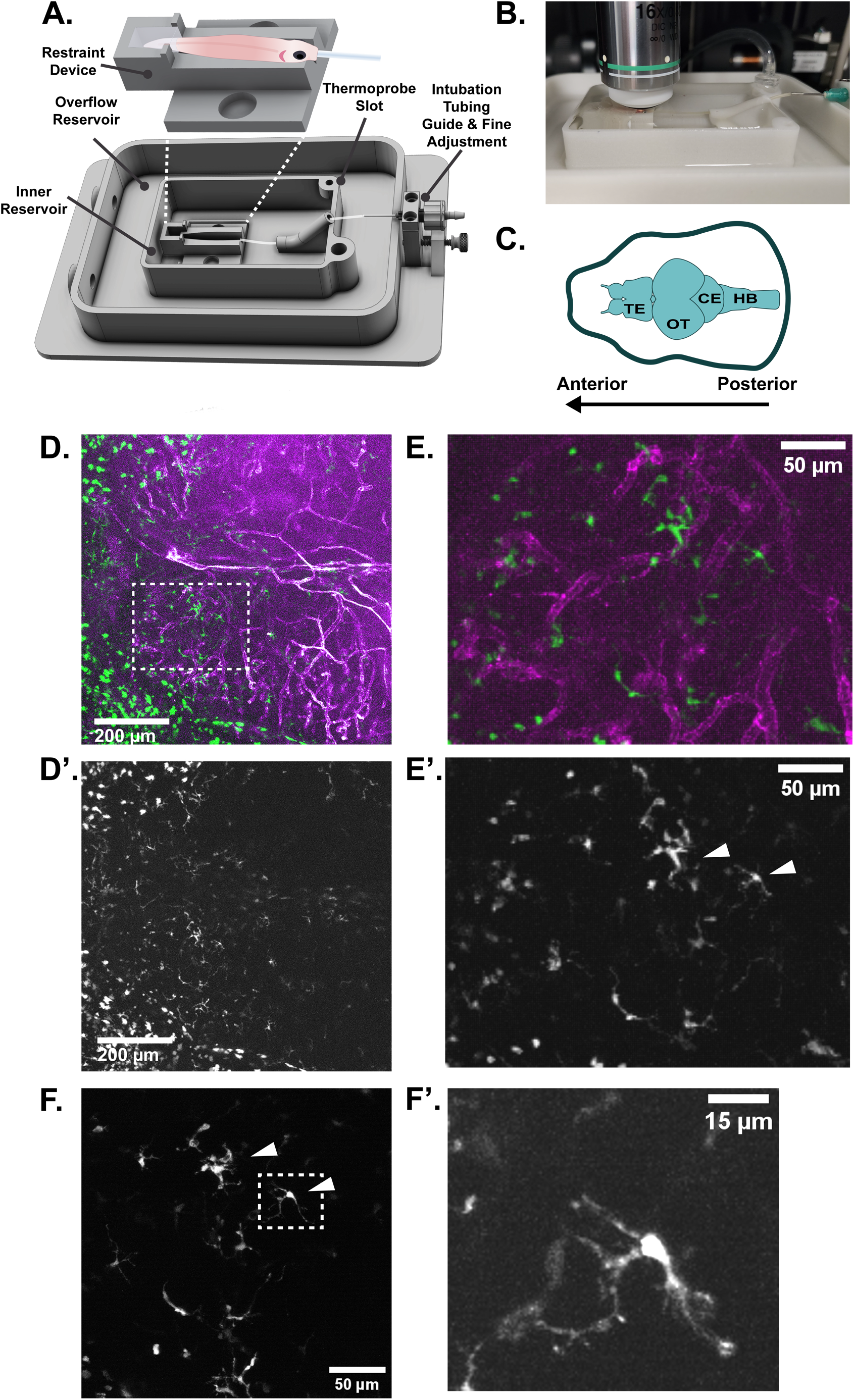
Non-invasive neuroimaging in the adult zebrafish. **A)** The life-support chamber is a modular design consisting of a base that docks into the microscope stage. The base includes an inner water reservoir and an overflow reservoir that allows water to drain and recirculate. Temperature of inner reservoir water can be continuously monitored via a thermoprobe. The fish is mounted in a restraint device by fixation of the tail with low melting point agarose. The dimensions of the restraint device can be modified to accommodate multiple ages and sizes of fish. **B)** An adult fish mounted in life-support chamber ready to be imaged. **C)** A diagram of regions of the adult zebrafish brain. Imaging data is presented anterior facing left. TE = telencephalon, OT = optic tectum, CE = cerebellum, HB = hind brain. **D)** A maximum intensity projection of both lobes of the telencephalon of the adult zebrafish imaged by multiphoton microscopy. Magenta = vasculature, as marked by mCherry driven by the *kdrl* promotor. Green = neuroimmune cells marked by eGFP driven by *mpeg1* promotor. **D’)** The eGFP channel from D isolated to better show neuroimmune cells. **E)** An enlargement of the inset marked by the white dotted line in D. Cells of ramified and amoeboid morphology are present. **E’)** The eGFP channel from E isolated to better show neuroimmune cells. White arrows indicate cells imaged in F. **F)** A still from a movie taken at 3× zoom showing a subset of cells present in E’ (white arrows). **F’)** A still of the individual cell demarcated by the white dashed box in F showing the amount of cellular detail that can be captured in live intravital imaging.

In agreement with previous intravital imaging studies of microglia, we found that ramified *mpeg1*:eGFP positive cells in young adult animals are highly dynamic in their protrusive behaviors (Fig. 2, Supplemental Movie 2). We recorded no significant translocation of the cell bodies of these cells during the imaging window. We also found that total cell volume did not change dramatically but instead was balanced between cycles of protrusion and retraction. This was consistent between cells in the telencephalon (Fig. 2A) and the optic tectum (Fig. 2B). These results confirm that adequate resolution for the study of the cell biology and dynamic behavior of neuroimmune cells can be achieved with non-invasive 2-photon imaging. It is likely that different cell types, such as other glia and neurons, could also be successfully imaged non-invasively using this method.

**Figure 2.**
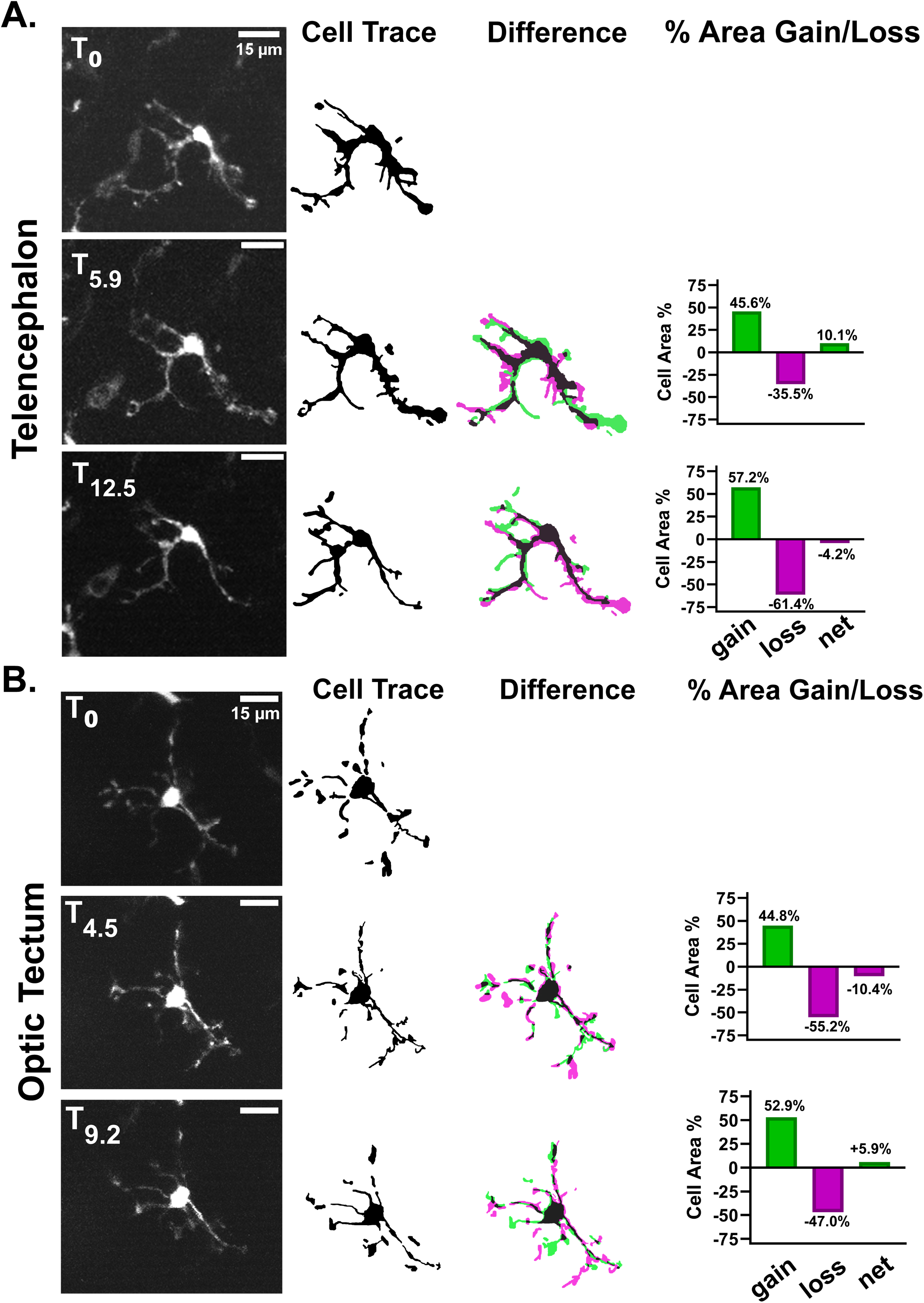
Intravital imaging allows monitoring of highly dynamic processes in *mpeg1:eGFP*+ neuroimmune cells across brain regions. **A)** Representative images from approximately 13 minutes of timelapse intravital imaging from the same animal depicted in figure 1D. Cells were traced and the difference between depicted frames was determined by overlaying cell traces (middle panel). The area of the cell volume gained (green) or lost (magenta) between the represented frame and the previous frame was calculated and is represented in the graphs to the right. The same procedure was performed in **(B)** except the imaged cell was in the optic tectum. No significant translocation of cell bodies was detected in either brain region.

### Repeated imaging of the same brain region in the same individual over months

Since we are interested in following changes in neuroimmune cell morphology over time and during disease progression, we wanted to confirm that we could reproducibly image the same brain region serially in the same individual. We tested if longitudinal imaging could be achieved non-invasively, without skin removal or skull thinning. We imaged a series of individual animals, including the same individual from Figures 1 and 2, every two weeks for 6 weeks in both the telencephalon and optic tectum (Fig. 3). Animals were imaged for approximately 2 hours each session. Rough estimates of imaging area were obtained through transmitted light visualization, then adjusted based on imaging of the vasculature and autofluorescence of the skull and meninges. The vasculature could be used successfully to identify specific regions and reliably re-identify them over weeks (Fig. 3C). Local neuroimmune cells were more difficult to reliably re-identify, as morphology and location changed significantly over the scale of the intervals between imaging sessions.

**Figure 3.**
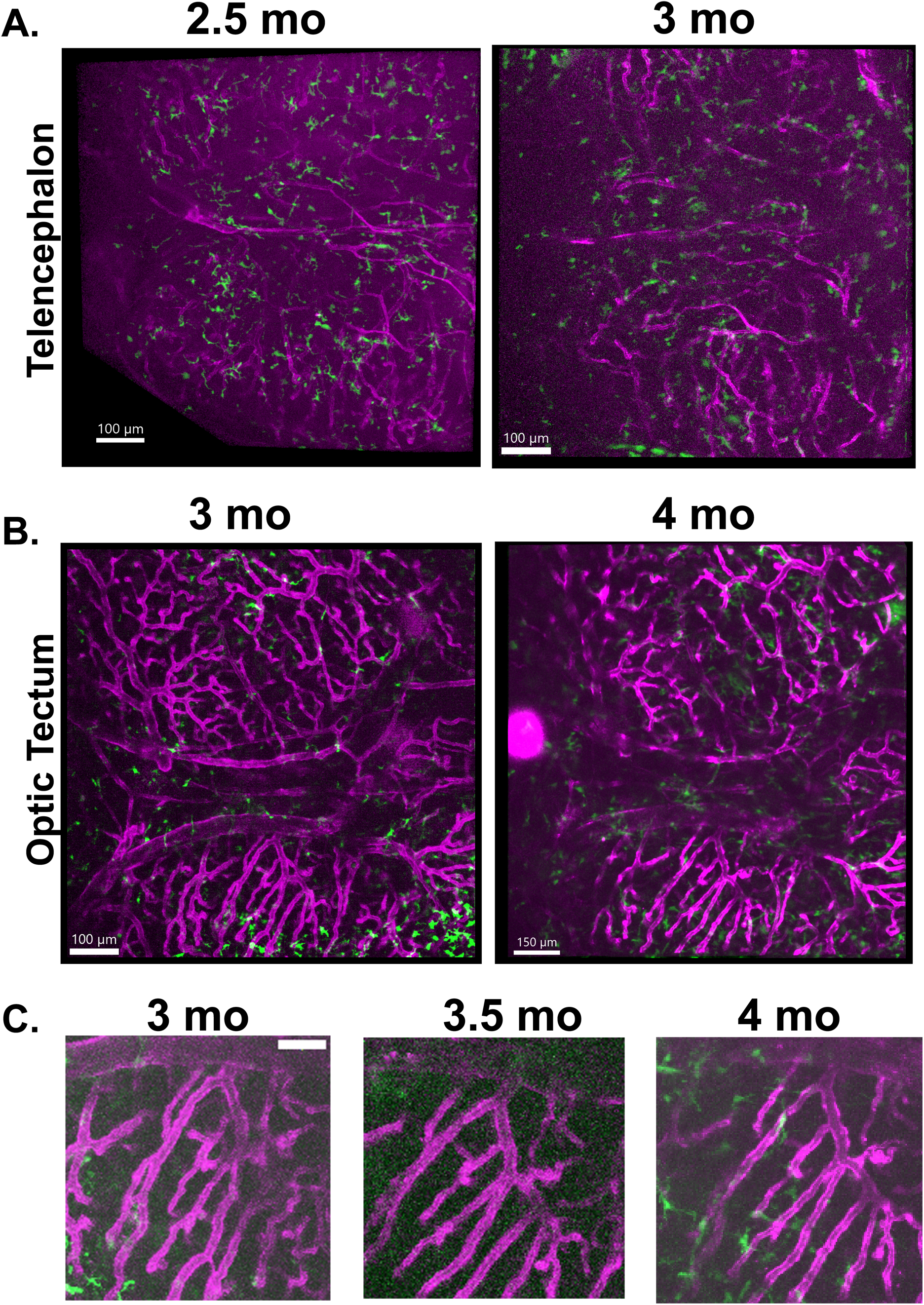
Longitudinal multiphoton imaging can reliably follow changes in the same brain region over weeks and months. The same animal as depicted in figures 1 and 2 was imaged across several weeks. Imaris was used to make projections of brain regions with skull and skin layers removed. **A)** The telencephalon of an individual animal with transgenic labeling of vasculature (*kdrl:mCherry*, magenta) and phagocytic neuroimmune cells (*mpeg1:eGFP*, green) was imaged at 2.5 months and again at 3 months. The vasculature can be compared to confirm similar areas. The morphology of the neuroimmune cells changes significantly over weeks, in agreement with previous data showing the dynamicity of their processes. **B)** A region of the optic tectum is imaged in the same individual animal at 3 months and again at 4 months old. **C)** Regions of the vasculature from the optic tectum of the same animal imaged at 3 months, 3.5 months, and 4 months old demonstrating that the same region can be reliably identified across imaging periods.

### Observing changes in neuroimmune cell morphology and behavior across the lifespan

Finally, we tested whether much older and larger animals were suitable for non-invasive intravital imaging. We were able to image animals ranging from 1-2 years old and obtain a similar degree of detail and ability to capture dynamic cellular behavior (Fig. 4, Supplemental Movies 3,4). No skull thinning or tissue removal was performed to obtain these results. Imaging older fish does pose some additional challenges, as we observed the distance between the skull and the first layers of brain tissue becomes larger. There is also an increase in structures we believe to be subcutaneous fat pockets that can have a distorting effect. The best imaging outcomes can be obtained by using brightfield imaging to identify areas with minimal fat pockets before imaging. We found that both the telencephalon and optic tectum remained accessible for imaging in these older animals. We were able to image a similar field of view for the telencephalon in aged adult animals as in younger animals (Fig. 4 A). Specific sub-regions of the telencephalon were identified (Fig. 4 A’, A’’) and selected for live imaging for periods of up to 30 minutes (Supplemental Movie 4, 5). Phagocytic neuroimmune cells within the brains of aged zebrafish displayed increased dynamic behavior and motility compared to that of younger animals. We were able to observe examples of cell-cell interaction (Fig. 4 B) and retraction of cellular process (Fig. 4 C) during our imaging window. Unlike in young animals, where little translocation of cell bodies was observed, tracked cells in aged animals showed increased cell motility as measured by average velocity (Fig. 4D).

**Figure 4.**
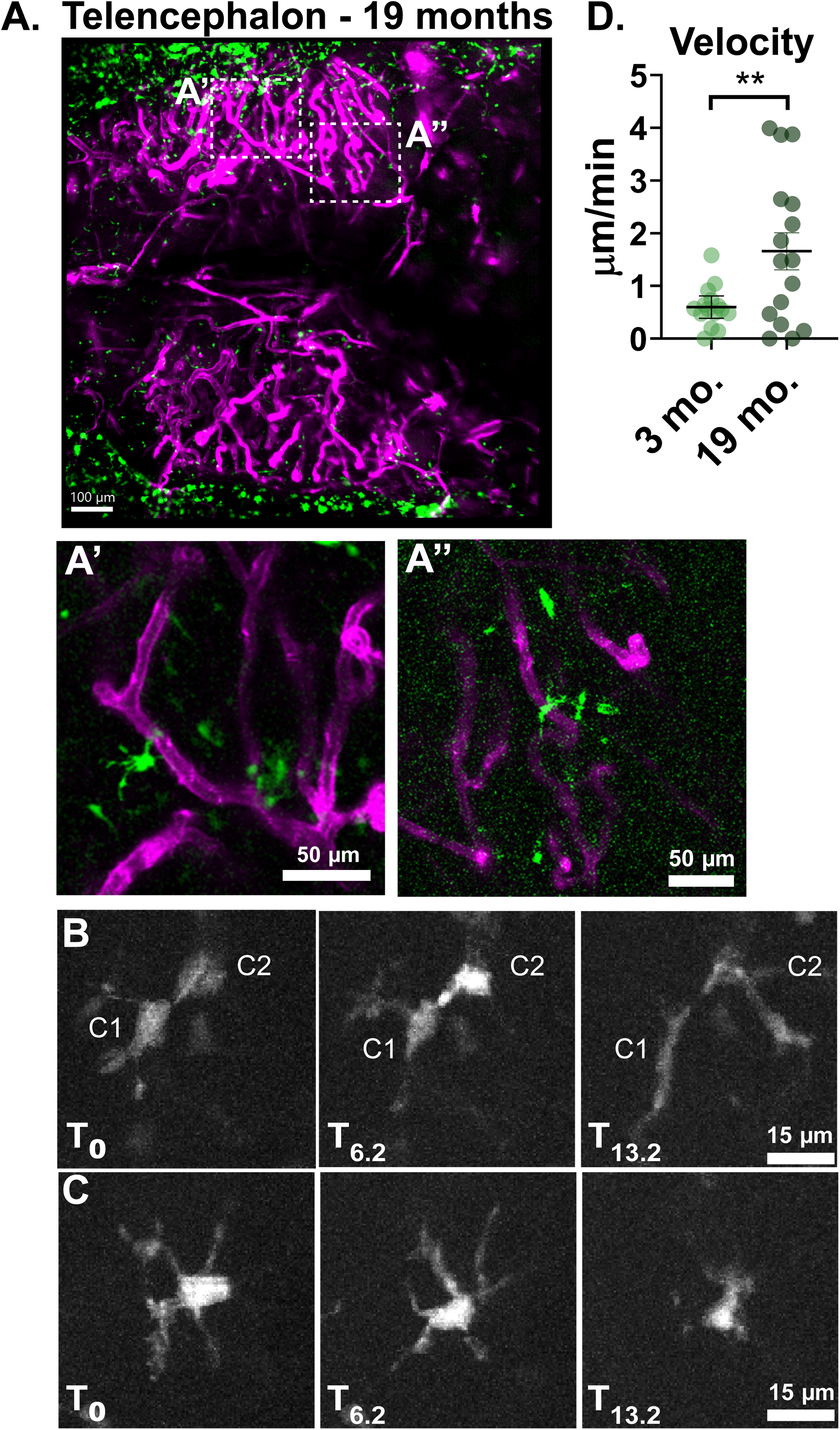
Longitudinal intravital imaging can detect changes in neuroimmune cell behavior between young and aged animals. **A)** The telencephalon of a 19-month old zebrafish with vasculature in magenta and neuroimmune phagocytes in green. **A’, A’’** are enlarged insets of the regions marked by white dashed boxes in A. **B-C)** Representative stills from a 14 minute long movie of the region depicted in A’. **B** shows two individual cells (C1, C2) touching and then elongating in response to contact. **C** shows an individual cell that retracts its protrusions over the course of the imaging period. **D)** To demonstrate that these methods allow us to detect differences in neuroimmune cell dynamics across ages, cells in two sample movies from a 3-month old animal were tracked and compared to cells from two sample movies from a 19-month old animal. Tracks were corrected for drift. Cells in the 19-month old animal demonstrated greater velocity and distance translocated than cells in the 3-month old animal. Velocity was calculated by summing the distance traveled per frame per cell, subtracting the drift, and then dividing the distance traveled by the imaging period. P=0.0099, unpaired T-test with Welch’s correction. Points represent individual cells.

## Discussion

We have developed and tested a chamber intended for live intravital neuroimaging of adult zebrafish. Zebrafish are a cost-effective and efficient model organism to conduct longitudinal non-invasive imaging for the study of neuroimmune cell biology. We have demonstrated that it is possible to non-invasively image the superficial layers of two regions of the zebrafish brain, the telencephalon and optic tectum, longitudinally across the lifespan of the zebrafish. We are particularly interested in non-invasive imaging since we are studying the cell biology of the neuroimmune system, where creating wounds or environmental disturbances can complicate interpretation of imaging data. However, in cases where this is not a concern, improved imaging depth could likely be achieved through previously demonstrated techniques like scale removal or skull thinning (Barbosa *et al*., 2016). We have shown that non-invasive imaging is suitable for imaging eGFP expressing cells up to 200μm deep, however the use of brighter and more red-shifted fluorophores or 3-photon illumination could further improve depth of imaging. Additionally, photoconvertible fluorophores may be necessary to accurately identify individual cells in local populations longitudinally. Further study is needed to compare the behavior and morphology of *mpeg1*:eGFP positive neuroimmune cells in superficial layers versus deeper layers. This technique could be expanded for imaging of other neuronal cell types, and the modular design of the restraint device design means it could be easily modified and redesigned to accommodate different mounting orientations for imaging different structures.

Finally, we have also shown preliminary evidence that like mammals, neuroimmune cells in zebrafish undergo detectable changes in morphology and behavior during aging. These changes in morphology and behavior are similar to observed morphological changes associated with microglial dysfunction in mammals (Hefendehl *et al*., 2014). While intriguing, further research is necessary to identify whether all neuroimmune populations across zebrafish brain regions show similar signs of aging, or if there are regional differences. Better transgenic lines with expression more restricted to mark microglia are also necessary to confirm the identity of specific neuroimmune populations. Since zebrafish are well known for their lifelong neuroproliferative ability, these preliminary findings pose many compelling questions about how neuroproliferative regions may influence neuroimmune aging. Though there is abundant work remaining to be done, we believe this device and these data strengthen and broaden the utility of the adult zebrafish model as a tool for neuroimaging and neurobiological study.

## Materials and Methods

### Chamber Design and Manufacture

We designed a device to facilitate fast and reproducible mounting of adult zebrafish fish with minimal restraint. This device can be 3D printed in biocompatible resin, Nylon-12, or produced in glass-filled nylon (preferable for long-term use). The design features a removable **restraint device** (Fig. 1 A, B, Sup. Fig. 2, 3) which secures the fish in place via a V-shaped channel and a small well for low-melting point agarose. The use of the well confines the agarose to only the tail of the fish and allows the fish to be freed from the chamber with minimal effort. Importantly, this prevents low melting point agarose from contacting the fish’s gills. Dental wax (widely available from drugstores) must be affixed over top of the fish’s back for further stabilization (Fig. 1B, Sup. Fig. 3 A’’)

The restraint device docks securely via magnets into the **inner reservoir** (Fig. 1A, Sup. Fig. 2G). We have designed the reservoir to include a second **overflow reservoir** (Fig. 1A), and two water outlet channels to prevent overflows. For unattended imaging, we recommend adding water overflow sensors (Castranova *et al*., 2022). The inner reservoir also features a dock for a temperature monitor probe (Fig. 1A). The fish is intubated during the imaging period to allow for adequate oxygenation and delivery of anesthetic. To increase the ease of the intubation process, we have added a thumbscrew-adjustment on the intubation tubing support (Fig. 1A, Sup. Fig. 2C) to change the position of the tubing quickly and accurately. This allows for fine adjustments of tube position to accommodate different lengths/ages of fish.

This imaging device was designed in SolidWorks to fit an ASI stage insert and can easily be modified for other stages. CAD files were sent for 3D printing (Midwest Prototypes) in glass-filled Nylon 12 using laser sintering. Glass-filled Nylon 12 is both biocompatible and strong and rigid making it resistant to warping. It can also be easily sterilized via soaking in 10% bleach without degradation of the material. A complete diagram of assembly steps can be found in Supplemental Figure 2, and additional assembly instructions are included in supplemental methods and online (https://morgridge.org/research/labs/fablab/designs/).

### Animals

Adult zebrafish (*Danio rerio*) were kept in a 14/10 hr. light/dark cycle and fed Gemma Micro 500. Serially imaged fish were separated into distinguishable pairs in 1L tanks on a quarantine rack to allow for accurate re-identification of individual animals. Transgenic lines Tg(*kdrl*:mCherry, *mpeg1*:eGFP)(Jin *et al*., 2005) and Tg(*mpeg1*:eGFP) (Ellett *et al*., 2011) were maintained in a pigmentless Casper roy^a9^, nacre^w1^ double mutant (Lister *et al*., 1999; Ren *et al*., 2002; White *et al*., 2008). We have found that nacre^w1^ mutants were also suitable for intravital imaging, and the presence of iridophores does not prevent imaging. Equal numbers of male and female animals were used. All animals in these studies were handled in accordance with the National Institutes of Health Guide for the care and use of laboratory animals, and the University of Wisconsin Institutional Animal Care and Use Committee (IACUC). These studies were approved by the University of Wisconsin IACUC (protocol L006536).

### Anesthesia

#### General considerations for anesthesia

As previously noted, multiple strategies have been used for anesthetizing adult zebrafish for microscopy. The goals for sustained anesthesia for long term microscopy differ from those of general anesthesia for brief procedures. MS-222 (tricaine) has been widely adopted for use in zebrafish for anesthesia. However, there are no standardized guidelines for adult dosage based on age and/or weight in zebrafish. We found that for animals over 60 days of age and above 180mg, maintenance concentrations of tricaine in the range of 160mg/L to 170mg/L is effective and well-tolerated (Supplemental Figure 1). Tricaine is tolerated very well in embryonic, larval, and adult zebrafish. However, we and others have found that the metamorphic stages in juveniles (roughly ages 21-70 days) tolerate tricaine poorly and have an increased risk of death during periods of anesthesia regardless of length of exposure or concentration of MS-222 (Owen and Kelsh, 2021). Therefore, we do not recommend imaging of fish in juvenile stages using MS-222 if it is vital that the fish be recovered. Alternative anesthetic strategies (such as combined use of isoflurane with tricaine) could also be considered (Huang *et al*., 2010; Lockwood *et al*., 2017; Owen and Kelsh, 2021). Use of a chamber that allows full encasement in agarose in place of tricaine to restrain the adolescent during imaging is also a viable strategy (Lam, 2022).

A major drawback to the use of tricaine as an anesthetic for neural imaging is that it can increase respiration (and thus gill movement) (Huang *et al*., 2010). This can result in increased jitter, drift, and imaging artifacts. At our current resolution, most jitter can be resolved computationally. However, in certain instances where it is crucial to have continuity of frame over time, paralytic agents such as pancuronium bromide or full encasement in low-melting point agarose could be considered. Both methods will likely affect the length of time the fish can be viably imaged, and the ability to recover the fish after imaging.

#### Anesthesia Protocol

Tricaine anesthetic (Syncaine by Syndel) was prepared at a stock concentration of 10mg/mL and gradually adjusted to pH 7.0 with Tris buffer (pH 9.0). Tricaine is highly acidic when dissolved in water so the pH should be carefully controlled to ensure comfortable anesthesia. If tricaine is not dissolved correctly, it can have an appearance like oil droplets on the surface. This can happen if pH is brought up too quickly, or not properly dissolved. We recommend discarding the solution if this occurs and consider a lower stock concentration if continued problems occur. Water for initial induction is prepared in a 1L tank at a concentration of 180-200mg/L in fresh system water. The maintenance dose of tricaine is prepared in the main reservoir at a concentration range of 150-170mg/L. Fish were fasted for at least 15 hours prior to imaging to avoid regurgitation. Adult fish were anesthetized by placement into the induction tank until loss of righting response was observed and the fish becomes unresponsive to a gentle tail pinch. After induction was achieved the fish remained deeply anesthetized for several minutes, during which mounting and intubation can be performed. If intubation is not achieved within ∼3-5 minutes, the fish may begin to show signs of movement and will have to be re-induced. The goal is to maintain a very light anesthesia for maximum chance of routine survival. Body standard length has been previously reported as an indicator of anesthesia tolerance in embryonic and larval zebrafish (Owen and Kelsh, 2021), but length becomes more variable in adults older than 90 days and is not reliable. We assessed anesthesia dosage per body weight and found animals weighing 180mg and above to be the most reliable for anesthesia at tricaine concentrations of 160-170mg/L (Sup. Fig. 1). Animals in the metamorphic (juvenile) state, where reliance on passive oxygen diffusion through the skin begins to wane in favor of active oxygen uptake via the gills, are more susceptible to tricaine overdose and exhibit unpredictable responses to tricaine(Owen and Kelsh, 2021). Therefore, we included only fully adult animals in this study. The age of the tricaine stock is also very important. Tricaine made from powder nearing the end of its shelf-life and frozen tricaine stocks nearing their expiration date have resulted in less predictable outcomes in our hands.

### Mounting

All materials were prepared and at hand prior to initial induction of anesthesia in the fish, since the time between induction and intubation should only be brief (2-3 minutes). Prior to starting the imaging device was securely placed on the microscope stage, the peristaltic pump and tubing were attached to the device, and water was warmed to 28° C and circulating in the chamber.

Low melting point (LMP) agarose (2% in PBS, Sigma-Aldritch) is prepared in aliquots, melted at 75° C and maintained at 42° C in a heat block in preparation for mounting. The restraint device is prepared by wrapping parafilm around the agarose reservoir that will secure the tail (Sup. Fig. 3A). After induction of anesthesia in the fish was confirmed (detailed in Anesthesia section) the fish was removed from the induction tank by using a pair of soft tipped tweezers to grip the tail fin and immediately placed into the groove of the restraint device (Sup. Fig. 3A). The device was tilted from horizontal to vertical and softly tapped on the counter to allow the fish’s tail to slide back into the agarose reservoir (Sup. Fig. 3B). If the tail was caught on top of the parafilm, soft-tipped forceps or a pipette tip were used to position the tail into the agarose reservoir. The fish should easily fall into an upright position due to the device shape, but upright positioning can be ensured by gently using soft-tipped forceps to adjust. Once the position was correct, LMP agarose is immediately added to the reservoir area of the restraint device while it is still held vertically (Sup. Fig. 3B). Maintaining a mostly vertical position ensures that agarose was kept away from the gills. No agarose should be beyond the anal fin of the fish (Sup. Fig. 3A’, dotted line). The LMP agarose was allowed to solidify. An ice pack can be helpful to hasten this process as long as care is taken to ensure that the fish does not come in direct contact with the ice pack. Once the agarose is solidified, the chamber was immediately placed into the inner chamber containing water with Tricaine at the maintenance dosage. Dental wax was draped over the upper body of the fish and pinched onto the sides of the restraint device to further stabilize the fish for imaging (Sup. Fig. 3A’’). The fish was carefully intubated by placing the 0.5mm ID, 1mm OD silicone tubing into the fish’s mouth. Water flow from the intubation tube was kept at ∼6ml/min, corresponding to a pump rate of 26.5 rotations per minute with our specified pump and tubing. A more detailed protocol is available online at (https://morgridge.org/research/labs/fablab/designs/).

### Live Imaging and Recovery

All imaging was performed with a Bruker Ultima or Ultima 2P+ equipped with a Spectra Physics Insight ultrafast laser and a Nikon CFI75 LWD 16X water-dipping objective. The period between frames for live imaging varied based on stack size but typically ranged from 30-45 seconds for a total of 10-20 minutes per movie. Fish were imaged for a maximum of 2 hours each imaging session. Temperature at the chamber was maintained between 27° C and 28° C and was consistently monitored via a temperature probe. The imaging plane was identified by focusing past the skin and skull (identified by autofluorescent signal and appearance of skin macrophages) and to the point where characteristic brain blood vessels and sparse immune cells could be identified. After imaging was completed, the intubation tubing was removed, and animals were recovered from the chamber by unwrapping the parafilm and using the end of a forceps or wooden applicator stick to dislodge the agarose from the restraint device. The fish can then be easily slid out of the device into an individual tank containing fresh water for resuscitation. The agarose will generally fall off as soon as the fish resumes movement, though use of a transfer pipette to agitate the water around the fish’s tail can help this process. Fish were housed in as individuals or in distinguishable pairs and monitored for an hour after recovery. Fish were checked again 12 hours after imaging to ensure no unusual behavior or injuries were present. We have found that fish are quick to recover from light anesthesia, but if a fish does not begin to show signs of movement within a minute, a transfer pipette can be used to force water past the fish’s gills to increase oxygenation and speed recovery. All fish successfully recovered from imaging survived for over a year until they were euthanized according to normal stock maintenance practices in our animal protocol.

## Acknowledgements

We would like to acknowledge the useful discussions that have made this work possible: D. Castranova for the initial inspiration and valuable feedback to refine this imaging protocol, the Zebrafish Rock community for helpful feedback surrounding zebrafish anesthesia practices, and K. Stephenson and L. Krugner-Higby for assistance developing our animal protocol. This work would also not be possible without technical assistance from J. V. Chacko.

## Figure Legends

**Supplemental Figure 1.**
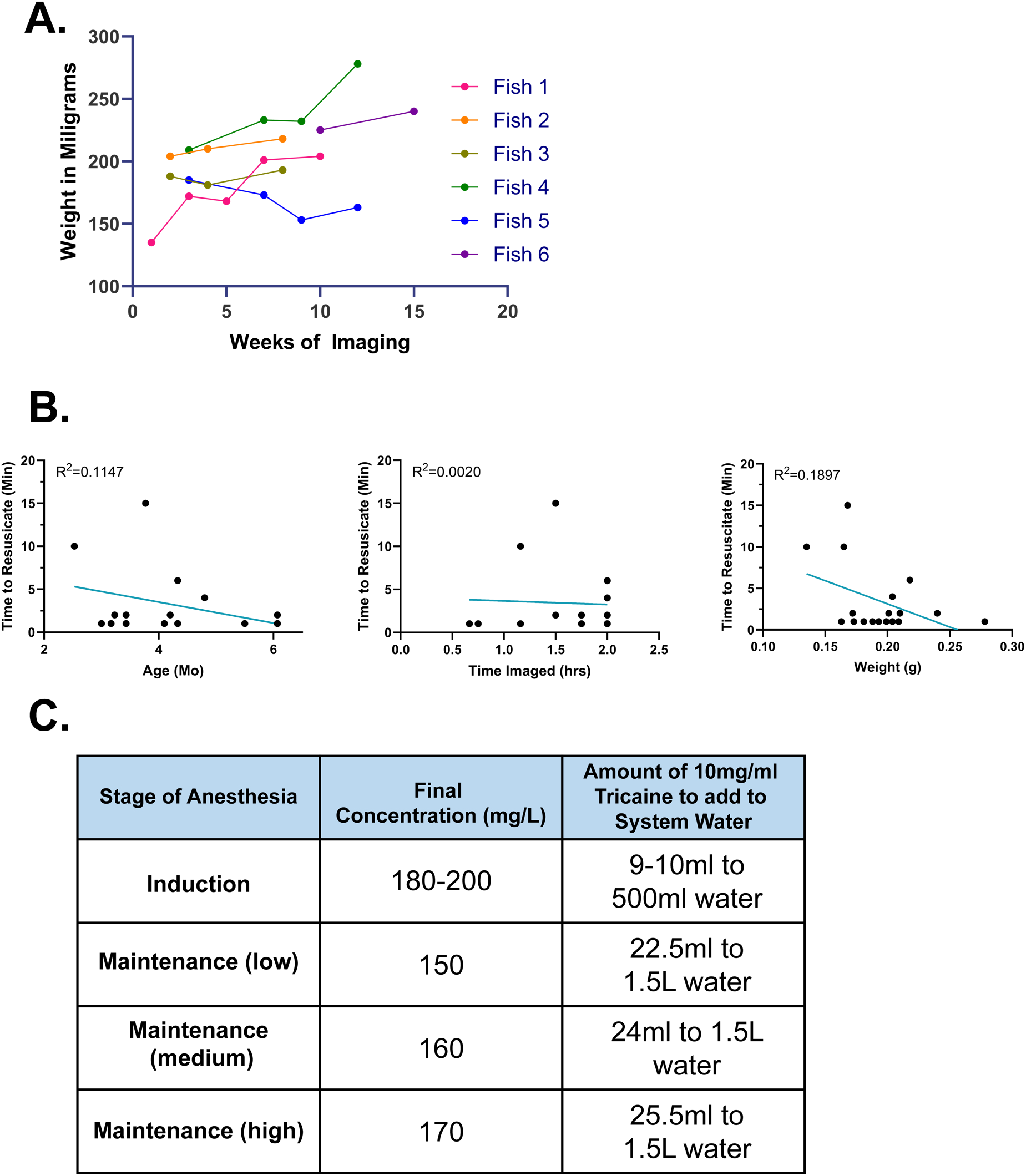
A) Fish subjected to longitudinal imaging did not demonstrate weight loss and continued to grow during the imaging period, suggesting that imaging did not interfere with normal feeding behavior or cause excessive stress. **B)** The time required to resuscitate an individual animal after imaging was not significantly correlated with any measured parameters. Time to Resuscitate vs Age: p = 0.115, slope = -1.22, Time to Resuscitate vs Time Imaged: p = 0.859, slope = -0.419, Time to Resuscitate vs Weight: p = 0.071, slope = 28.82. **C)** Table of tricaine dosage based on stage of anesthesia for fish older than 60 days old (past metamorphic stage).

**Supplemental Figure 2.**
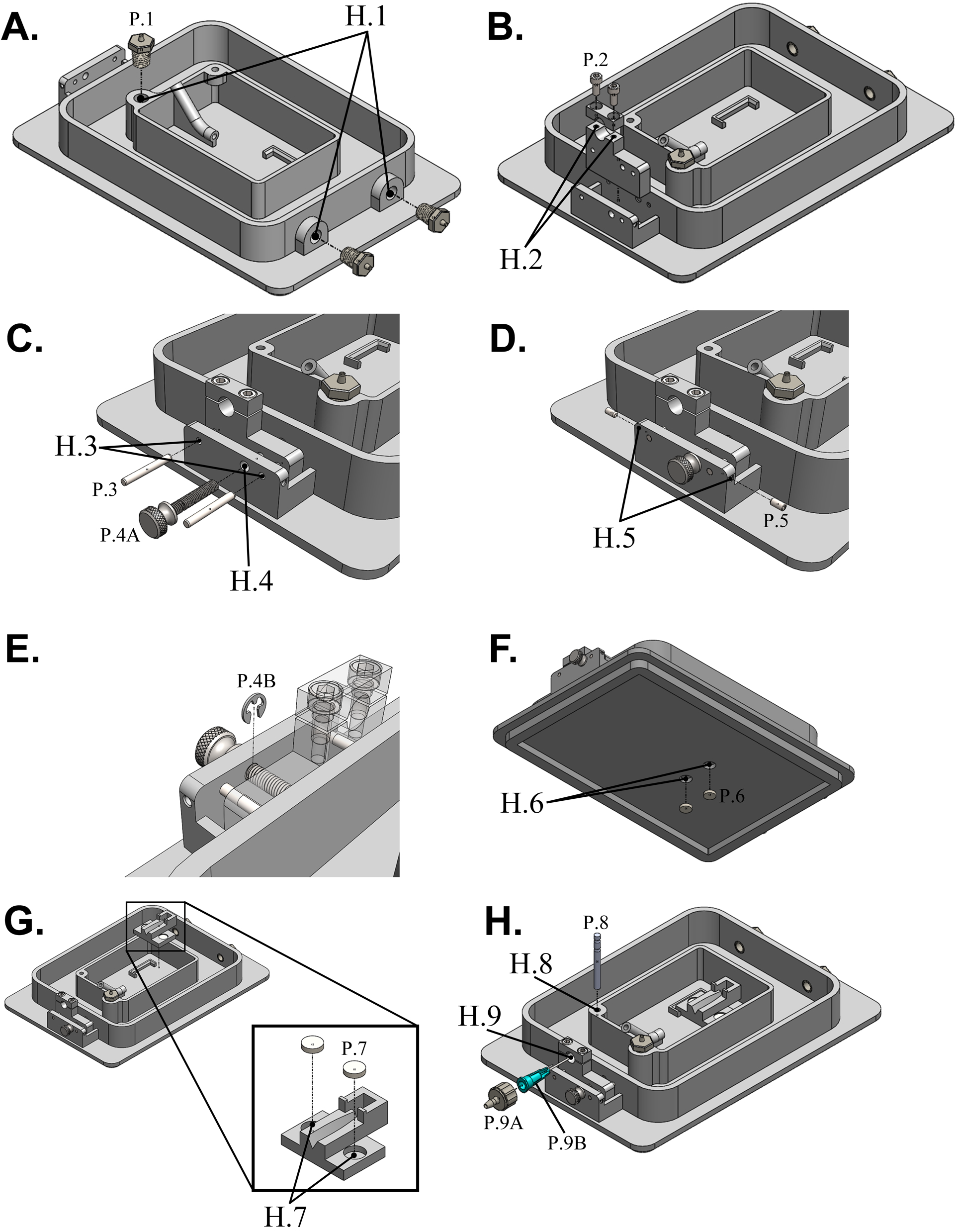
Chamber assembly and full feature diagram. **A)** The inflow hole and outflow holes (**H.1**) are shown. All holes are tapped using a 1/16-27 NPT standard tap and threading-to-barb fittings (1/16” ID tube) are screwed in. **B)** Holes for screws in the top of the support (H.2) are tapped using a #4-40 UNC tap. Socket screws are screwed in to tighten and loosen the clamp as needed. **C)** Steel dowel pins are placed in H.3, going through the outer chamber and the support piece. A thumb screw is placed in H.4, with only the part of the hole through the support piece being tapped. A M3x0.5 tap is used. The thumb screw must have a 0.5 mm groove starting 3.5 mm from the collar and heading away from the collar. The resulting diameter in the groove must be 3/32”. **D)** Set screws to secure the dowel pins are placed in the side holes (**H.5**). These side holes are tapped with a M2×0.4 tap. **E)** An external retaining ring is placed in the groove on the thumb screw to secure it in place. **F)** A small amount of epoxy is placed in H.6, then magnets are inserted. More epoxy is placed over the magnets to protect them from corrosion. **G)** Magnets are placed in H.7 in the inner chamber using the same method as the previous step. The magnets snap the inner chamber into place but still allow for easy removal. **H)** The digital thermometer probe is placed in the holder (**H.8**) in the inner walls and the blunt end needle and Luer lock adapter is secured in the support (**H.9**).

**Supplemental Figure 3.**
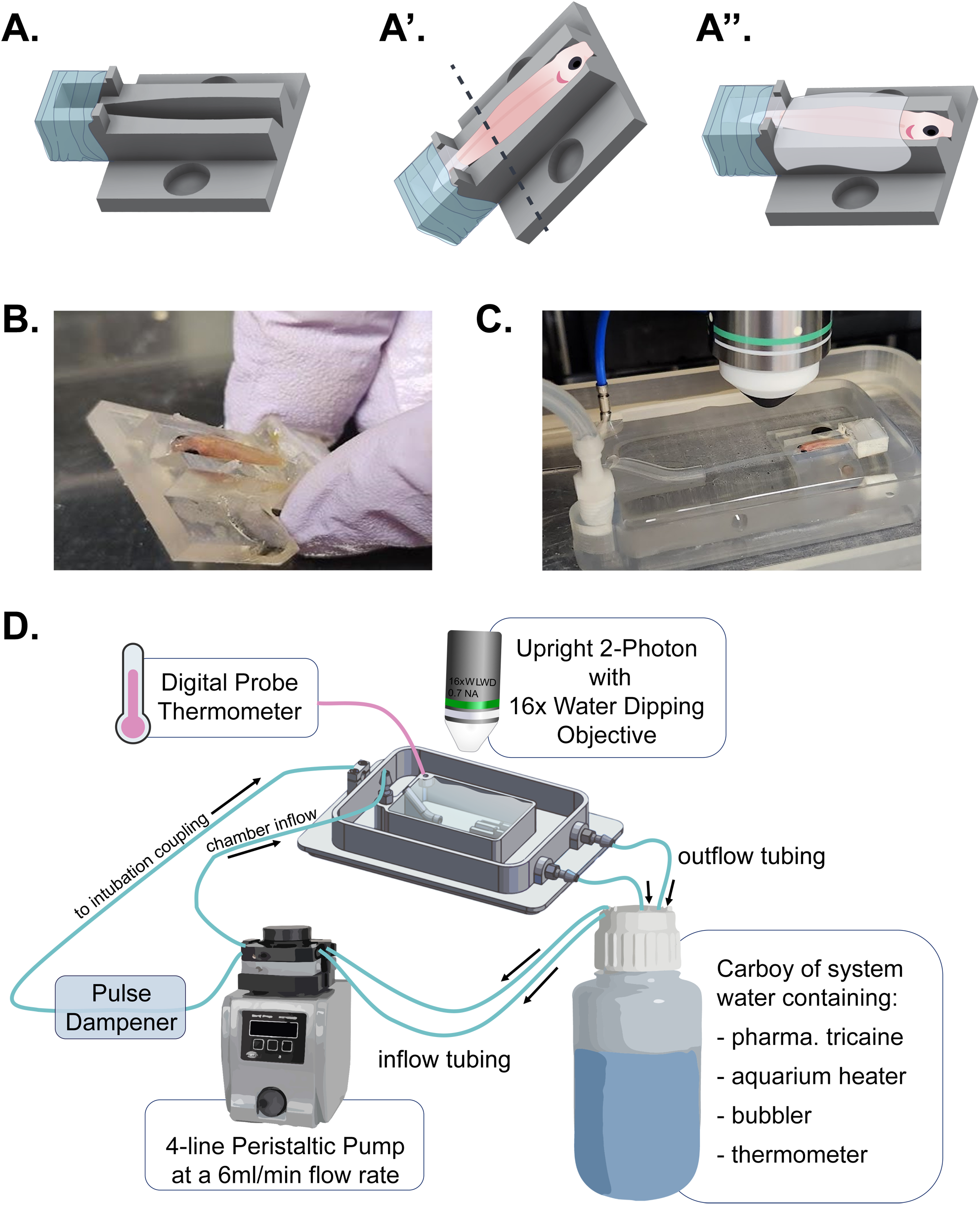
Mounting of fish and imaging setup. **A)** The restraint device is prepared by wrapping parafilm around the low melting point agarose reservoir. **A’)** The anesthetized fish is placed into the prepared restraint device and tilted upwards (tapping on the benchtop helps fish’s tail slide into the agarose reservoir). Low melting point agarose can be added to reservoir by pipetting and should not fill the reservoir past the dotted line to prevent interference with gills. Caution should be taken to ensure that agarose is an appropriate temperature to avoid burns to fish. **A’’)** A blanket of dental wax can be draped across the fish’s back to further immobilize, with care taken to leave head and gills exposed. Dental wax can be pinched to side of restraint device to secure. Once agarose is set, the device is ready to be placed into the inner reservoir of imaging chamber. **B)** Example image of fish in restraint device prior to adding dental wax. **C)** Fish placed in restraint device in imaging chamber without addition of dental wax. Intubation tube is placed in the fish’s mouth once it is inside chamber. Caution: take care not to position the intubation tube too deeply in the fish’s mouth to prevent damage or death to animal. **D)** Schematic of total intubation setup. A perfusion pump is used to circulate warmed system water containing tricaine anesthetic through a pulse dampener and into the imaging chamber. Water is continuously circulated and overflows the top of the inner chamber into the overflow chamber. Water outflows from the overflow chamber and returns to the main carboy where it can be rewarmed and oxygenated.

**Supplemental Movie 1**. An imaris rendering showing the superficial layers of skin and skull being computationally removed to show the lobes of the telencephalon of a young adult zebrafish imaged by multiphoton microscopy. Magenta is vasculature, as marked by mCherry driven by the *kdrl* promotor. Green is neuroimmune cells marked by eGFP driven by *mpeg1* promotor. Movie supports figure 1.

**Supplemental Movie 2**. Two movies depicting dynamic movements of *mpeg1* positive neuroimmune cells in the telencephalon (TE, top) and optic tectum (OT, bottom) of an adult zebrafish. Movie supports figures 1 and 2.

**Supplemental Movie 3**. An imaris rendering showing the superficial layers of skin and skull being computationally removed to show the lobes of the telencephalon of an aged adult zebrafish imaged by multiphoton microscopy. Magenta is vasculature, as marked by mCherry driven by the *kdrl* promotor. Green is neuroimmune cells marked by eGFP driven by *mpeg1* promotor. Movie supports figure 4.

**Supplemental Movie 4**. Two movies depicting dynamic movements of *mpeg1* positive neuroimmune cells in the telencephalon (TE, top) and optic tectum (OT, bottom) of an adult zebrafish. Movie supports figure 4.

